# The type IV pilin PilA couples surface attachment and cell cycle initiation in *Caulobacter crescentus*

**DOI:** 10.1101/766329

**Authors:** Luca Del Medico, Dario Cerletti, Matthias Christen, Beat Christen

## Abstract

Understanding how bacteria colonize surfaces and regulate cell cycle progression in response to cellular adhesion is of fundamental importance. Here, we used transposon sequencing in conjunction with FRET microscopy to uncover the molecular mechanism how surface sensing drives cell cycle initiation in *Caulobacter crescentus*. We identified the type IV pilin protein PilA as the primary signaling input that couples surface contact to cell cycle initiation via the second messenger c-di-GMP. Upon retraction of pili filaments, the monomeric pilin reservoir in the inner membrane is sensed by the 17 amino-acid transmembrane helix of PilA to activate the PleC-PleD two component signaling system, increase cellular c-di-GMP levels and signal the onset of the cell cycle. We termed the PilA signaling sequence CIP for cell cycle initiating pilin peptide. Addition of the chemically synthesized CIP peptide initiates cell cycle progression and simultaneously inhibits surface attachment. The broad conservation of the type IV pili and their importance in pathogens for host colonization suggests that CIP peptide mimetics offer new strategies to inhibit surface-sensing, prevent biofilm formation and control persistent infections.

**Significance Statement:** Pili are hair-like appendages found on the surface of many bacteria to promote adhesion. Here, we provide systems-level findings on a molecular signal transduction pathway that interlinks surface sensing with cell cycle initiation. We propose that surface attachment induces depolymerization of pili filaments. The concomitant increase in pilin sub-units within the inner membrane function as a stimulus to activate the second messenger c-di-GMP and trigger cell cycle initiation. Further-more, we show that the provision of a 17 amino acid synthetic peptide corresponding to the membrane portion of the pilin sub-unit mimics surface sensing, activates cell cycle initiation and inhibits surface attachment. Thus, synthetic peptide mimetics of pilin may represent new chemotypes to control biofilm formation and treat bacterial infections.

---

The cell cycle model bacterium *Caulobacter crescentus* (*Caulobacter* thereafter) integrates surface colonization into a bi-phasic life-cycle. Attachment begins with a reversible phase, mediated by surface structures such as pili and flagella, followed by a transition to irreversible attachment mediated by polysaccharides (1–4). In *Caulobacter* surface sensing is intimately interlinking with cellular differentiation and cell cycle progression (5, 6). During the bi-phasic life cycle, *Caulobacter* divides asymmetrically and produces two distinct cell types with specialized development programs (Fig. 1a). The sessile stalked cell immediately initiates a new round of chromosome replication, whereas the motile swarmer cell, equipped with a polar flagellum and polar pili, remains in the G1 phase for a defined interval before differentiating into a stalked cell and entering into the replicative S phase driven by the second messenger c-di-GMP dependent degradation of the cell cycle master regulator CtrA (7, 8) (Fig. 1a). The change in cell cycle state from motile swarmer into surface attached replication-competent stalked cells depends on tactile sensing mechanisms. Both pili and flagella have been previously implicated as key determinants involved in tactile surface sensing (9, 10). However, understanding the molecular mechanism of how *Caulobacter* interlinks bacterial surface attachment to cell cycle initiation has remained elusive.

**Fig. 1.**
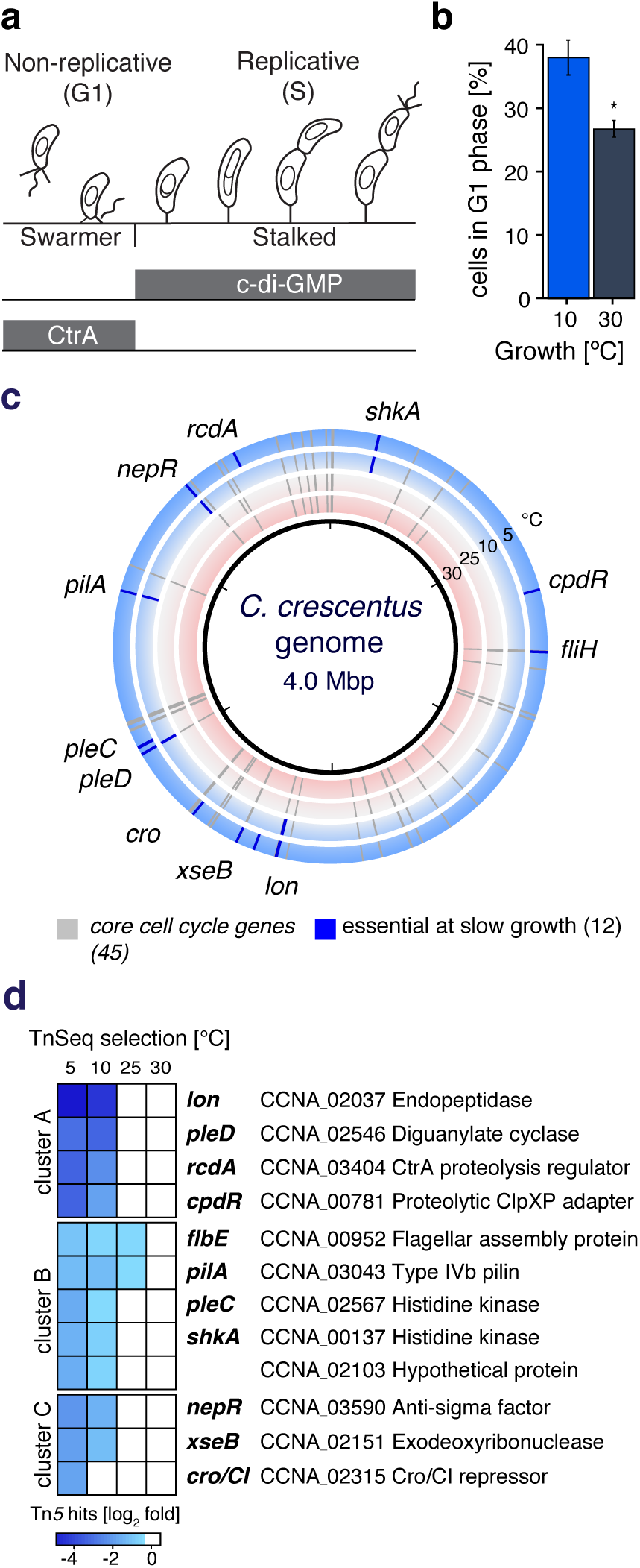
Transposon sequencing identifies conditionally essential genes required during reduced growth. **(a)** *Caulobacter* divides asymmetrically into a replication competent stalked cell and a swarmer cell. The master regulator CtrA inhibits DNA replication in swarmer cells and is proteolytically cleared upon an increase in cellular levels of the second messenger c-di-GMP at the G1-S phase transition. **(b)** Cellular replication under slow-growth conditions increases the relative duration of the G1-phase. **(c)** TnSeq across the 4.0 Mbp *Caulobacter* genome defines 45 core-essential cell cycle genes (grey marks) to sustain growth at 5°C, 10°C, 25°C and 30°C (outer to inner track) and 12 conditionally essential genes required for slow-growth conditions (blue marks). **(d)** Hierarchical cluster analysis of the 12 conditionally essential genes required for slow growth.

In this work, we report on a short peptide signal encoded within the type IVb pilin protein PilA that exerts pleiotropic control and links bacterial surface attachment to cell cycle initiation in *Caulobacter*. Using FRET microscopy in conjunction with a genetically encoded c-di-GMP biosensor (11), we quantify c-di-GMP signalling dynamics inside single cells and found that, besides its structural role in forming type IVb pili filaments, monomeric PilA in the inner membrane functions as a specific input signal that triggers c-di-GMP signalling at the G1-S phase transition.

## Results

### A specific cell cycle checkpoint delays cell cycle initiation

To understand how bacterial cells adjust the cell cycle to reduced growth conditions, we profiled the replication time of the α- proteobacterial cell cycle model organism *Caulobacter* across the temperature range encountered in its natural freshwater habitat (Table S1). Under the standard laboratory growth temperature of 30°C, *Caulobacter* replicates every 84 ± 1.2 min. However, when restricting the growth temperature to 10°C, we observed a 13-fold increase in the duration of the cell cycle, extending the replication time to 1092 ± 14.4 min (Table S1). To investigate whether reduced growth resulted in a uniform slow-down or affects particular cell cycle phases, we determined the relative length of the G1 phase by fluorescence microscopy using a previously described cell cycle reporter strain (11) (Materials and Methods). We found that the culturing of *Caulobacter* at 10°C caused a more than 1.4-fold increase in the relative duration of the G1 phase indicating a delay in cell cycle initiation (Fig. 1b). This finding suggested the presence of a specific cell cycle checkpoint that delays cell cycle initiation during reduced growth conditions.

### TnSeq identifies conditionally essential cell cycle genes

To identify the complete set of genes required for cell cycle progression at different growth rates, we designed a systems-wide forward genetic screen based on quantitative selection analysis coupled to transposon sequencing (TnSeq) (12, 13). TnSeq measures genome-wide changes in transposon insertion abundance upon subjecting large mutant populations to different selection regimes and enables genome-wide identification of essential genes. We hypothesised that the profiling of growth-rate dependent changes in gene essentiality will elucidate the components of the cell cycle machinery fundamental for cell cycle initiation under reduced growth conditions. We selected *Caulobacter* transposon mutant libraries for prolonged growth at low temperatures (5°C and 10°C) and under standard laboratory cultivation conditions (25°C and 30°C). Cumulatively, we mapped for each condition between 397’377 and 502’774 unique transposon insertion sites across the 4.0 Mbp *Caulobacter* genome corresponding to a transposon insertion densities of 4-5 bp (Table S2).

To identify the factors required for cell cycle progression, we focused our analysis on essential genes (Data SI) that are expressed in a cell cycle-dependent manner (Materials and Methods, (14–16)). Among 373 cell cycle-controlled genes, we found 45 genes that were essential under all growth conditions (Fig. 1c, Data SI), including five master regulators (*ctrA, gcrA, sciP, ccrM* and *DnaA*), eleven divisome and cell wall components (*ftsABILQYZ, fzlA* and *murDEF*), six DNA replication and segregation factors (*dnaB, ssb, gyrA, mipZ, parB* and *ftsK*) as well as 23 genes encoding for key signalling factors and cellular components required for cell cycle progression (Data SI). Collectively, these 45 genes form the core components of the bacterial cell cycle machinery.

### Components of the c-di-GMP signalling network are conditionally essential for slow growth

During reduced growth conditions, we found 12 genes that specifically became essential (Fig. 1c). To gain insights into the underlying genetic modules, we performed a hierarchical clustering analysis and grouped these 12 genes according to their growth-rate dependent fitness profile into three functional clusters A, B and C (Fig. 1d, Materials and Methods).

Cluster A contained four conditionally essential genes that exhibited a large decrease in fitness during slow-growth conditions (Fig 1d, Fig. S1). Among them were *pleD, cpdR, rcdA* and *lon* that all comprise important regulators for cell cycle controlled proteolysis. The diguanylate cyclase PleD produces the bacterial second messenger c-di-GMP, which becomes restricted to the staked cell progeny upon asymmetric cell division and is absent in the newly born swarmer cell (Fig. 1a, (17)). During the G1-S phase transition, c-di-GMP levels raise again and trigger proteolytic clearance of the cell cycle master regulator CtrA (Fig 1a), which is mediated by ClpXP and the proteolytic adaptor proteins RcdA, CpdR and PopA (18–20). Similarly, the ATP-dependent endopeptidase Lon is responsible for the degradation of the cell cycle master regulators CcrM, DnaA and SciP (21–23). Taken together, these findings underscore the importance to control proteolysis of CtrA and other cell cycle regulators to maintain cell cycle progression at low growth rates when intrinsic protein turnover rates are marginal.

Cluster B contained five genes including the two kinase genes *pleC* and *shkA*, a gene of unknown function encoding for a conserved hypothetical protein (CCNA_02103) as well as the type IV pilin gene *pilA* and the flagellar assembly ATPase *flbE/fliH* (Fig 1d, Fig. S2). Multiple genes of cluster B participate in c-di-GMP signalling. Among them, we found the sensor kinase PleC that functions upstream and activates the diguanylate cyclase PleD over phosphorylation. The hybrid kinase ShkA comprises a downstream effector protein of PleD, which binds c-di-GMP and phosphorylates the TacA transcription factor responsible for the initiation of the stalked-cell specific transcription program (24). The flagellum assembly ATPase FlbE/FliH together with FliI and FliJ form the soluble component of the flagellar export apparatus, which in *Pseudomonas* has also been identified as a c-di-GMP effector complex (25). Collectively, these data indicate that c-di-GMP signalling is of fundamental importance to coordinate cell cycle progression under slow-growth conditions.

Cluster C comprised the anti-sigma factor *nepR*, a Cro/CI transcription factor (CCNA_02315) and *xseB* encoding the small subunit of the exodeoxyribonuclease VII (Fig 1d, Fig. S3), which form outer-circle components not directly linked to c-di-GMP signalling. However, disruption of these genes likely induces stress response and alternative transcriptional programs that may interfere with cell cycle progression under slow-growth conditions.

### PilA induce c-di-GMP signalling

Among the components of the c-di-GMP signalling network identified within cluster A and B, the sensor kinase PleC resides at the top of the signalling hierarchy activating the diguanylate cyclase PleD at the G1-S phase transition. While the activation mechanism of its down stream target PleD has been resolved with molecular detail (26), the type of external signal integrated by PleC has remained unknown. PleC comprises an amino-terminal periplasmic domain, which suggests that PleC activity is controlled by an external signal. Among the identified conditionally essential gene within cluster B, PilA co-clustered together with PleC (Fig 1d). The Type IV pilin protein PilA is translocated into the periplasm and shares the same sub-cellular localization pattern to the swarmer specific cell pole as PleC (17, 27). Thus, we speculated that PilA is a likely candidate for the hitherto unknown external input signal perceived by the sensor kinase PleC.

To test this hypothesis, we monitored the c-di-GMP signalling dynamics in a wildtype and Δ*pilA* background. The activation of PleC, and subsequently PleD, leads to a strong increase in the c-di-GMP levels at the G1-S phase transition (Fig. 1a, (11)). Any mutation that impairs activation of PleC is expected to prolong the G1-swarmer phase. To monitor c-di-GMP signalling dynamics inside single cells, we have previously engineered a genetically encoded FRET-biosensor that permits time-resolved monitoring of the fluctuating c-di-GMP levels along the cell cycle (17). We synchronized wildtype and Δ*pilA* cells expressing this FRET biosensor and quantified c-di-GMP levels in individual swarmer cells by FRET-microscopy (Fig. 2a, Materials and Methods). In the wildtype control, we observed that the majority of synchronized cells quickly transitioned into the S phase with only 11.8% (143 out of 1073 cells) remaining in the G1 phase as indicated by low c-di-GMP levels (Fig. 2a,b). In contrast, we observed that more than 52.2% (821 out of 1570 cells) of all synchronized Δ*pilA* mutant cells exhibited a delay in the G1-S transition and maintained low c-di-GMP levels for a prolonged interval (Fig. 2a,b). Similarly, using time-lapse studies to follow signalling trajectories of individual cells, we found that Δ*pilA* mutants exhibited a more than 1.8 fold increase in the duration of the G1 phase as compared to wildtype cells (Fig. 2c). Providing a episomal copy of *pilA* restored the cell cycle timing defect of a Δ*pilA* mutant strain (Fig. 2a,b). These findings establish the type IV pilin PilA as a novel cell cycle input signal.

**Fig. 2.**
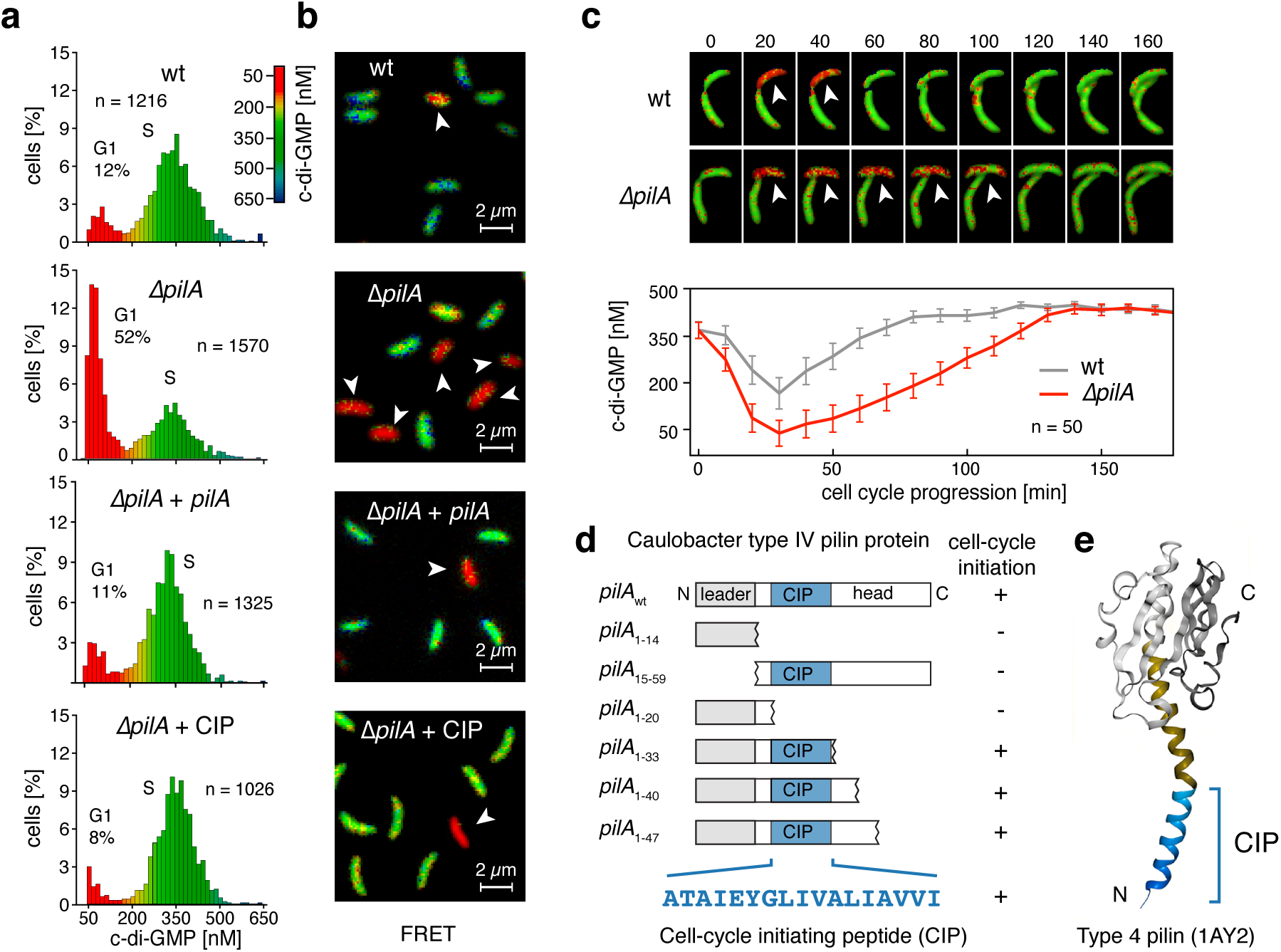
The CIP sequence encoded in the N-terminal portion of PilA functions as cell cycle initiating signal. **(a)** Population distribution of intra-cellular c-di-GMP concentrations in synchronized *Caulobacter* cells. The population shows a bimodal c-di-GMP distribution corresponding to swarmer cells (G1) prior and after (S) the G1 to S transition. **(b)** (Top) Dual-emission ratio microscopic (FRET) images of synchronized swarmer populations of wild-type *Caulobacter*, Δ*pilA* mutants Δ*pilA* complemented by a plasmid born copy of *pilA* and Δ*pilA* complemented by the exogenous addition of cell cycle initiating peptide CIP (100 uM). Pseudocolors show FRET emission ratios (527/480 nm) corresponding to the cytoplasmic c-di-GMP concentration as indicated by the color bar. Swarmer cells (highlighted by arrows) resting in the G1 phase prior initiation of cell cycle exhibit low cellular c-di-GMP levels. **(c)** Kinetics of fluorescence ratio changes (527/480 nm), reflecting c-di-GMP levels recorded in *Caulobacter* cells during the S to G1 transition. Upper panel: Time-lapse dual-emission ratiometric FRET microscopy of representative cells of *Caulobacter* wild-type (wt) and Δ*pilA* mutants recorded at intervals of 10 min. Lower panel: Corresponding plots of the measured c-di-GMP fluctuations for the indicated strains over time. The average single cell FRET ratio over a population of 50 cells upon cell division is shown for wild type (grey) and Δ*pilA* (red). The drop in c-di-GMP levels is larger and sustained much longer for the Δ*pilA* mutant. **(d)** Complementation of the cell cycle initiation defect of Δ*pilA* with a panel of N- and C-terminal truncated PilA variants. **(e)** The CIP peptide sequence is modeled onto the type IV pilin from *Neisseria* that harbours a larger globular C-terminal domain (grey) which is absent in the *Caulobacter* PilA protein (gold).

### PilA controls cell cycle initiation via the sensor kinase PleC and the diguanylate cyclase PleD

PleC is a bifunctional phosphatase/kinase that switches activity in a cell cycle dependent manner (28). In the newborn swarmer cell, PleC first assumes phosphatase activity to inactivate the diguanylate cyclase PleD and establish low cellular c-di-GMP levels. Subsequently, PleC switches to a kinase and activates PleD at the G1-S-phase transition. To test whether PilA functions as an input signal for the PleC-PleD signalling cascade, we measured c-di-GMP signalling dynamics and compared the duration of the G1 phase in single and double deletion mutants using FRET microscopy (Materials and Methods, Fig. S4). Similar to Δ*pilA* mutants, Δ*pleD* mutants also prolonged the G1 phase more than two-fold as compared to the wildtype control (18% and 22% versus 9% G1 cells in wildtype, Fig. S4). However, a Δ*pilA*, Δ*pleD* double mutant only marginally increased the duration of the G1 phase and showed similar frequencies of swarmer cells with low-c-di-GMP levels as compared to a strain lacking solely *pleD* (28% and 22% G1 cells, Fig S4). This genetic evidence suggest that PilA resides upstream of PleD. Thus, besides its role as a structural component of the type IVb pilus, PilA likely comprises an input signal for the sensor kinase PleC, which serves as the cognate kinase of PleD, to increase c-di-GMP levels at the G1-S phase transition.

### The inner membrane reservoir of the type IV pilin PilA signals cell cycle initiation

PilA harbours a short 14 aa N-terminal leader sequence required for translocation across the inner membrane that is cleaved off by the peptidase CpaA (29–31). To test whether translocation of PilA is a prerequisite for signalling the cell cycle initiation, we constructed a cytosolic version of PilA (*pilA*15-59) lacking the N-terminal leader sequence needed for the translocation of the matured PilA across the membrane. Unlike the full-length PilA, the episomal expression of the translocation-deficient version of PilA did not complement for the cell cycle timing defect of a Δ*pilA* mutant. Similarly, we found that solely expressing the N-terminal leader sequence of PilA (*pilA*1-14) neither restored the cell cycle timing defects of a Δ*pilA* mutant (Fig. 2d, Fig. S5, Table S3). We concluded that the translocation of PilA into the periplasm is a prerequisite to signal cell cycle initiation.

Upon translocation and cleavage of the N-terminal leader sequence, the mature form of PilA resides as a monomeric protein in the inner membrane (32). Through the action of a dedicated type IV pilus assembly machinery, the inner membrane-bound reservoir of PilA polymerizes into polar pili filaments that mediate initial attachment to surfaces (33). However, only *pilA* but none of the other component of the pilus assembly machinery became essential in our TnSeq screen under slow-growth conditions (Fig. 1c, Table S4, Data SI). Furthermore, unlike Δ*pilA* mutants, deletion mutants of the pilus assembly machinery genes *cpaA, cpaD*, and *cpaE* did not prolong the G1 phase but, in contrast, shorted the duration of the G1-phase two-fold as compared to the wildtype control (4%, 3%, and 5% G1 cells, Fig S4). The observation that the translocation of PilA monomers across the inner membrane is necessary but the subsequent polymerization of PilA monomers into mature pilin filaments is dispensable for cell cycle initiation, suggested that the periplasmic membrane reservoir of the monomeric form of PilA functions as an input signal for c-di-GMP mediated cell cycle signalling.

### The trans-membrane helix of PilA comprises a 17 amino-acid peptide signal that mediates cell cycle initiation

The mature form of PilA is a small 45 aa protein comprised of a highly hydrophobic N-terminal alpha-helix (α1N), which anchors PilA in the inner membrane (30, 32), and an adjacent variable alpha-helical domain protruding into the periplasm. To identify the portion of the matured PilA protein responsible for triggering cell cycle initiation, we constructed a panel of C-terminally truncated *pilA* variants and assessed their ability to complement the cell cycle defect of a chromosomal Δ*pilA* mutant by quantifying c-di-GMP dynamics in single cells using FRET-microscopy (Materials and Methods). The expression of a truncated PilA variant that includes the leader sequence and the first 5 N-terminal amino-acids of the matured PilA protein (*pilA*1-20) did not complement the cell cycle initiation defect of a Δ*pilA* mutant (Fig. 2d, Fig. S5, Table S3). However, increasing the N-terminal portion of the matured PilA protein to 17, 25, and 37 amino acids (*pilA*1-33, *pilA*1-40, *pilA*1-47) restored the cell cycle initiation defects of a Δ*pilA* mutant (Fig. 2d, Fig. S5, Table S3). Based on these findings, we concluded that a small N-terminal peptide sequence covering only 17 N-terminal amino acids from the mature PilA (Fig. 2e) is sufficient to initiate c-di-GMP dependent cell cycle progression. Accordingly, we annotated these 17 amino acids as <u>c</u>ell cycle initiating <u>p</u>ilin sequence (CIP) (Fig. 2e).

### Chemically synthesized CIP peptide initiates cell cycle progression

Next, we asked whether the translocation of PilA from the cytoplasm into the inner membrane or the presence of a PilA reservoir in the periplasm is sensed. If signalling depends solely on the presence of membrane inserted PilA, we speculated that exogenous provision of chemical-synthesized CIP peptide should restore the cell cycle defects in a Δ*pilA* mutant. To test this hypothesis, we incubated synchronized swarmer cells of a Δ*pilA* mutant in the presence of 100µM of chemically synthesized CIP peptide and assayed c-di-GMP signalling dynamics by FRET microscopy (Fig. 2a,b). Indeed, we found that addition of the CIP peptide induced cell cycle transition into the S-phase as indicated by a 6.5 fold lower abundance of G1-swarmer cells with low c-di-GMP levels as compared to an untreated Δ*pilA* cell population (8% vs 52% G1 cells, Fig. 2a,b). These findings suggest that the addition of the CIP peptide shortens the G1 phase and, thus, functions as a cell cycle activator. Collectively, these result demonstrated that neither the translocation or polymerization but solely the inner membrane reservoir of PilA is sensed via the hydrophobic CIP sequence to initiate cell cycle progression.

### The CIP peptide reduces surface attachment and ΦCbk phage susceptibility

Deletion of the sensor kinase *pleC* results in pleiotrophic defects and causes daughter cells to omit the G1-phase with low c-di-GMP levels (Fig. S4, (17)). Furthermore, *pleC* deletion mutants lack polar pili and show defects in initial attachment to surfaces (34). The observation that the addition of the CIP peptide shortens the G1-phase, suggests that the CIP sequence of PilA modulates PleC activity to promote phosphorylation of the downstream effector PleD. To test this hypothesis, we investigated whether the addition of the chemically synthesized CIP peptide also impairs additional PleC-specific output functions. Indeed, when assaying for the presence of functional pili using pili-specific bacteriophage CbK, we found that the incubation of synchronized wildtype *Caulobacter* for 10 min with the CIP peptide resulted in a 77.2 ± 2.1% decrease in bacteriophage CbK susceptibility (Fig. 3a,b), suggesting that CIP impairs pili function. Furthermore, we also found that the addition of the CIP peptide to wildtype *Caulobacter CB15* cells reduced initial attachment by 71.3 ± 1.6% (Fig. 3c,d) with a half-effective peptide concentration (EC50) of 8.9 µM (Fig. S6) and a Hill coefficient of 1.6 suggesting positive cooperativity in binding of CIP to a multimeric receptor complex. Altogether, these findings support a model in which the CIP sequence of PilA functions as a pleiotropic small peptide modulator of the sensor kinase PleC leading to premature cell cycle initiation, retraction of type IV pili and impairment of surfaces attachment.

**Fig. 3.**
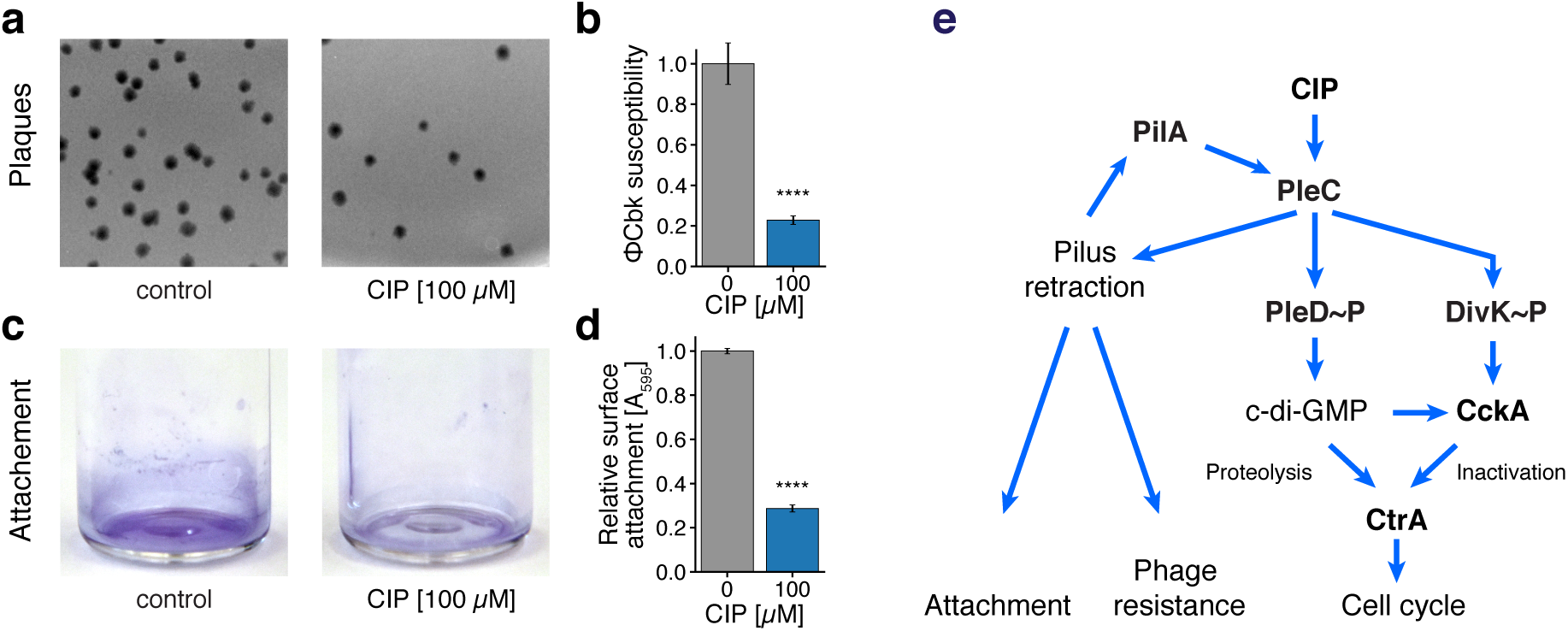
Pleotropic effects and proposed mode of action of the cell cycle initiating pilin peptide (CIP). **(a)** ΦCbk phage susceptibility assay of *Caulobacter* CB15 in the presence (right panel) and absence (left panel) of 100 µM CIP as detected by the formation of plaques on double agar overlay (n= 5). **(b)** Corresponding bargraph plot of the relative ΦCbk phage susceptibility of *Caulobacter* CB15 upon 10 min of incubation without (n = 5) or with (n = 5) 100 µM CIP peptide. Error bars = s.e.m., ****: P << 10-4. **(c)** Crystal violet surface attachment assay of *Caulobacter* CB15 in the presence (right panel) and absence (left panel) of 100 µM CIP. **(d)** Corresponding bargraph plot of the relative surface attachment of *Caulobacter* CB15 upon 1 h of incubation without (n = 9) or with (n = 9) 100 µM CIP peptide. Error bars = s.e.m., ****: P << 10-4. **(e)** Model for the mode of action of the CIP peptide. The kinase activity of the pleiotropic, membrane bound sensor kinase PleC is activated by the CIP peptide mimicking a inner membrane reservoir of PilA monomers. The CIP peptide modulates PleC activity to promote phosphorylation of the downstream effectors PleD and DivK as well as activates pili retraction as part of a positive feedback loop. CtrA activity must be removed from cells at the onset of DNA replication, because phosphorylated CtrA binds to and silences the origin of replication. The c-di-GMP and CckA signalling cascade orchestrates cell cycle entry through controlled proteolysis and inactivation of the master cell cycle regulator CtrA.

## Discussion

Understanding how bacteria regulate cell cycle progression in response to external signalling cues is of fundamental importance. In this study, we used a transposon sequencing approach to identify genes required for cell cycle initiation. Comparing deviations in gene essentiality between growth at low temperatures (5°C and 10°C) and under standard laboratory cultivation conditions (25°C and 30°C) allowed us to pinpoint genes required exclusively for cell cycle initiation. We identified the pilin protein PilA together with 6 additional components of a multi-layered c-di-GMP signalling network (Fig. 1) as key determinants that control cell cycle initiation. Using FRET microscopy studies, we quantified c-di-GMP signalling dynamics inside single cells and found that, besides its structural role in forming type IVb pili filaments, PilA comprises a specific input signal for activation of c-di-GMP signalling at the G1-S phase transition. Furthermore, we show genetic evidence that PilA functions upstream of the PleC-PleD two-component signalling system and present data that the monomeric PilA reservoir is sensed through a short 17 amino acid long N-terminal peptide sequence (<u>c</u>ell cycle initiation <u>p</u>ilin, CIP). It is remarkable that the *Caulobacter* PilA is a multi-functional protein that encodes within a 59 amino acid polypeptide a leader sequences for translocation, the CIP sequence for cell cycle signalling functions as well as structural determinants required for pili polymerization.

*Caulobacter* exhibits a biphasic life cycle where new-born swarmer progeny undergo an obligate differentiate into surface attached stalked cells to initiate DNA replication. How could PilA couple cell cycle initiation to surface sensing? Type IVb pili have been shown to serve as adhesion filament as well as mechano-sensors (35–37), that upon surface contact induce the retraction of pili filaments with concomitant accumulation of monomeric PilA in the inner membrane (9). Our results from genetic dissection of PilA in conjunction with FRET microscopy to monitor c-di-GMP signalling dynamics suggest a model where high levels of PilA monomers in the inner membrane are sensed via the N-terminal CIP sequence to activate the sensor kinase PleC and promote phosphorylation of the diguanylate cyclase PleD (Fig. 3e). In such a model surface sensing by pili induces a sharp increase in intra-cellular c-di-GMP levels and activates the proteolytic clearance (18, 20) of the master regulator CtrA to initiate cell cycle progression. However, the precise structural mechanism how initial surface contact stimulates pili retraction remains an open question. In *Caulobacter*, studies showed that pili upon covalent attachment of high-molecular-weight conjugates loose their dynamic activities (9). One possibility is that upon surface contact force generation leads to structural rearrangements within the filament that locks pili in the retraction state, which in turn prevents pilus extension and reincorporation of sub-units leading to elevated PilA levels in the inner membrane.

On the level of signal integration, sensing the PilA reservoir through the bifunctional kinase/phosphatase PleC provides robust signal integration to sense surfaces. Small fluctuations in the inner membrane PilA concentrations due to dynamic pili cycling in the planktonic state do not lead to permanent increase in cellular c-di-GMP levels as PleC kinase activity induced upon retraction is reversed when pili are extended again and PleC is reset to a phosphatase. In a model where surface contact locks pili in the retraction state, the kinase activity of PleC wins the tug of war leading to a permanent increase in c-di-GMP levels and a robust surface sensing mechanism.

From an engineering perspective, coupling surface sensing via the inner membrane PilA reservoir to a c-di-GMP second-messenger cascade may provide a mechanism to couple cell cycle initiation to multiple orthogonal output functions including inhibition of flagellar motility (38), secretion of adhesive surface polysaccharides (9) and activation of stalk-cell specific transcriptional programs (24) to induce permanent surface attachment and cellular differentiation at the G1-S phase transition.

The bacterial second messenger cyclic diguanylate (c-di-GMP) is a key regulator of cellular motility, cell cycle initiation, and biofilm formation with its resultant antibiotic tolerance, which can make chronic infections difficult to treat (39–41). In our work, we show that the addition of a chemically produced CIP peptide specifically modulates the c-di-GMP signalling behaviour in cells and also has pleiotropic effects on surface adhesion and phage susceptibility in *Caulobacter*. Therefore, the CIP peptide, regulating the spatiotemporal production of c-di-GMP, might be an attractive drug target for the control of biofilm formation that is part of chronic infections. Given the broad conservation of type IV pili and their central role in human pathogens to drive infection and host colonization (42), the CIP peptide identified represents a new chemotype and is potentially developable into a chemical genetic tool to dissect c-di-GMP signalling networks and to block surface sensing in pathogens to treat bacterial infections.

## Materials and Methods

Supplementary Materials and Methods include detailed descriptions of strains, media and standard growth conditions, Tn5 transposon mutagenesis, growth selection and sequencing, FRET microscopy procedures to quantify cellular c-di-GMP levels in *Caulobacter* cells, CIP peptide assay, quantification of surface attachment and phage susceptibility assays. Data SI contains the essentiality classification of each *Caulobacter* coding sequence profiled at different growth temperatures according to TnSeq measurements.

### TnSeq library generation

Tn*5* hyper-saturated transposon mutant libraries in *Caulobacter* were generated as previously described (12, 13). Transposon mutant libraries were selected on rich medium (PYE) supplemented with gentamicin. Depending on the respective library, the plates were incubated at the selection temperatures of 5, 10, 25 or 30°C. As soon as colonies appeared, the selected mutant libraries were separately pooled off the plates, supplemented with 10% v/v DMSO and stored in a 96-well format at -80°C for subsequent use.

### FRET microscopy

FRET imaging was performed on a Nikon Eclipse Ti-E inverted microscope with a precisExcite CoolLED light source, a Hamamatsu ORCA-ERA CCD camera, a Plan Apo λ 100x Oil Ph3 DM objective, combined with a heating unit to maintain an environmental temperature of 25°C during the imaging. Single time point acquisitions were taken under the acquisition and channel settings according to (11).

### CIP Peptide Assay

The CIP peptide was ordered from Thermo Fisher Scientific GENEART (Regensburg, Germany). Stocks were kept in 100% DMSO and the peptide was applied in a final DMSO concentration of 4% to synchronized *Caulobacter* NA1000 wt and Δ*pilA* populations prior FRET-microscopy. The swarmer fraction was resuspended in M2 salts and the CIP peptide was administered to a final concentration of 100 uM.

## ACKNOWLEDGMENTS

We thank C. Aquino and R. Schlap- bach from the Zurich Functional Genomics Center for sequencing support, W-D. Hardt, S.I. Miller for helpful comments on the manuscript. This work received institutional support from the Swiss Federal Institute of Technology (ETH) Zürich, ETH research grant [ETH-08 16-1] to B.C, and the Swiss National Science Foundation, [31003A_166476, 310030_184664 and CRSII5_177164] to B.C.

No conflict of interest declared.

---

**Fig. S1.**
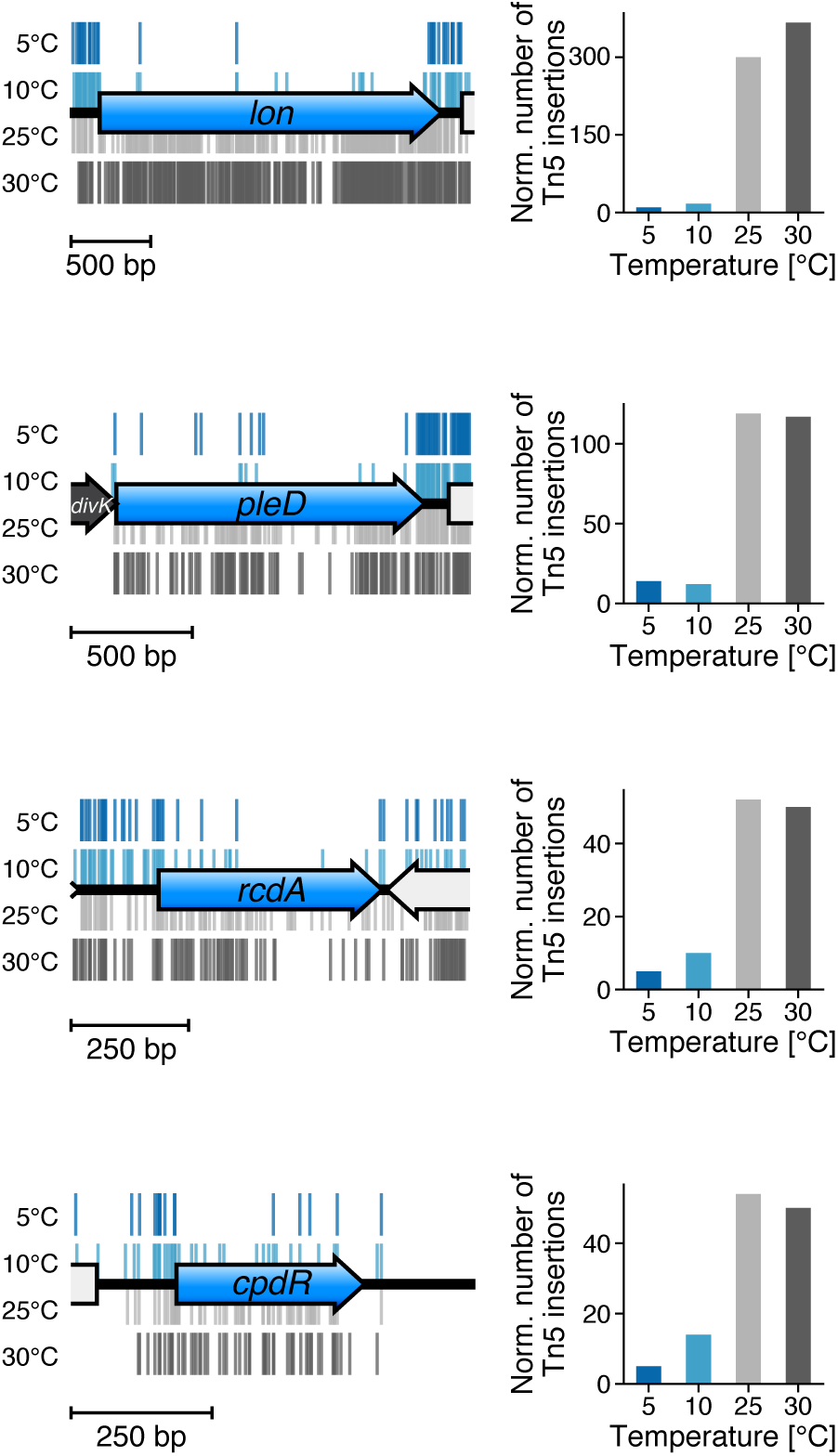
Tn*5* insertion patterns and numbers of insertions in conditional essential cell cycle genes of cluster A required for slow growth (blue arrows). The genomic positions of transposon insertions recovered upon selection a the respective growth temperature are plotted above and below the genome track as blue to dark grey marks. The normalized number of insertions within the open reading frame are plotted on the right.

**Fig. S2.**
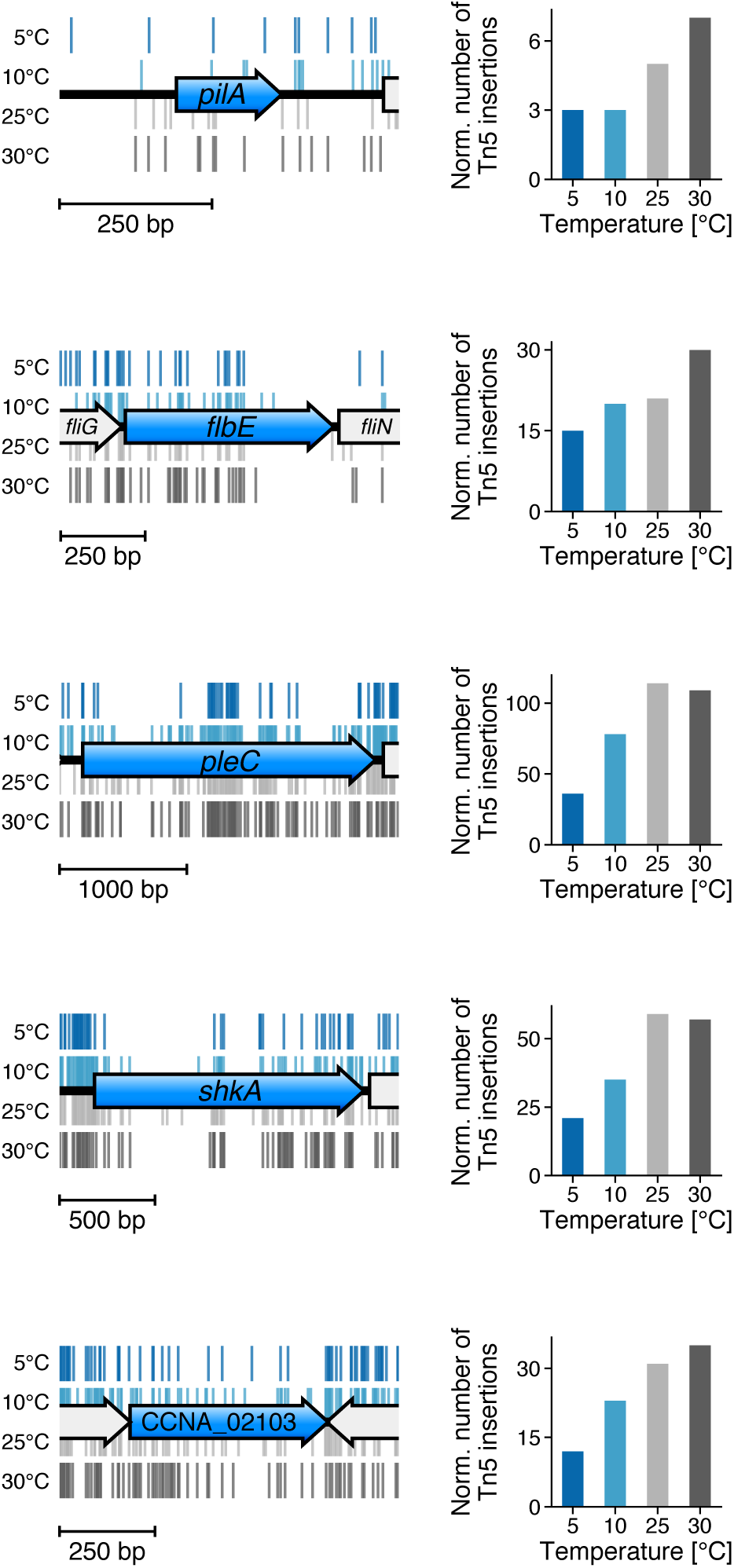
Tn*5* insertion patterns and numbers of insertions in conditional essential cell cycle genes of cluster B required for slow growth (blue arrows). The genomic positions of transposon insertions recovered upon selection a the respective growth temperature are plotted above and below the genome track as blue to dark grey marks. The normalized number of insertions within the open reading frame are plotted on the right.

**Fig. S3.**
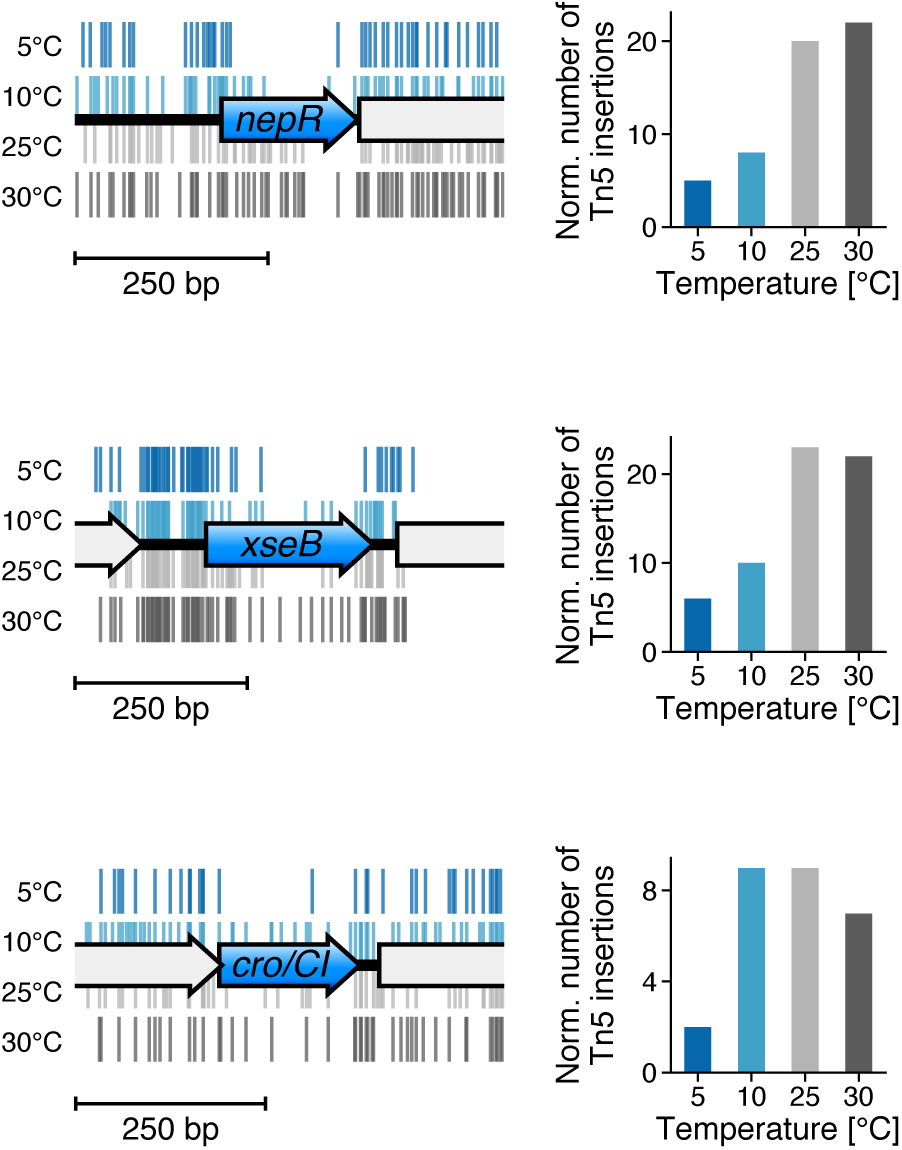
Tn*5* insertion patterns and numbers of insertions in conditional essential cell cycle genes of cluster C required for slow growth (blue arrows). The genomic positions of transposon insertions recovered upon selection a the respective growth temperature are plotted above and below the genome track as blue to dark grey marks. The normalized number of insertions within the open reading frame are plotted on the right.

**Fig. S4.**
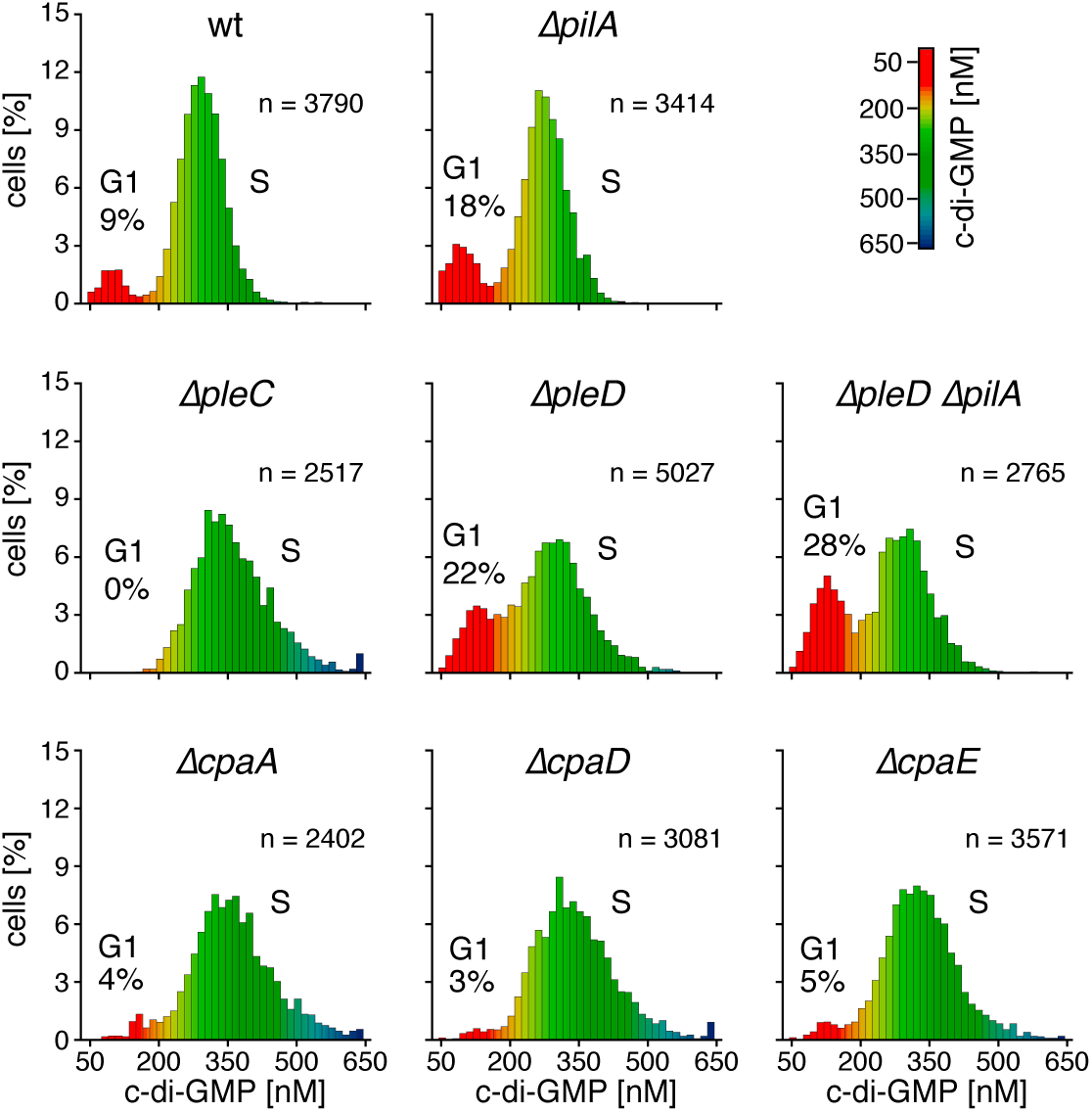
Population distribution of intra-cellular c-di-GMP concentrations in *Caulobacter* cells assessed by time-lapse FRET microscopy. Cells with low c-di-GMP levels correspond to swarmer cells (G1) while cells with high c-di-GMP concentrations correspond to replication competent stalked cells (S).

**Fig. S5.**
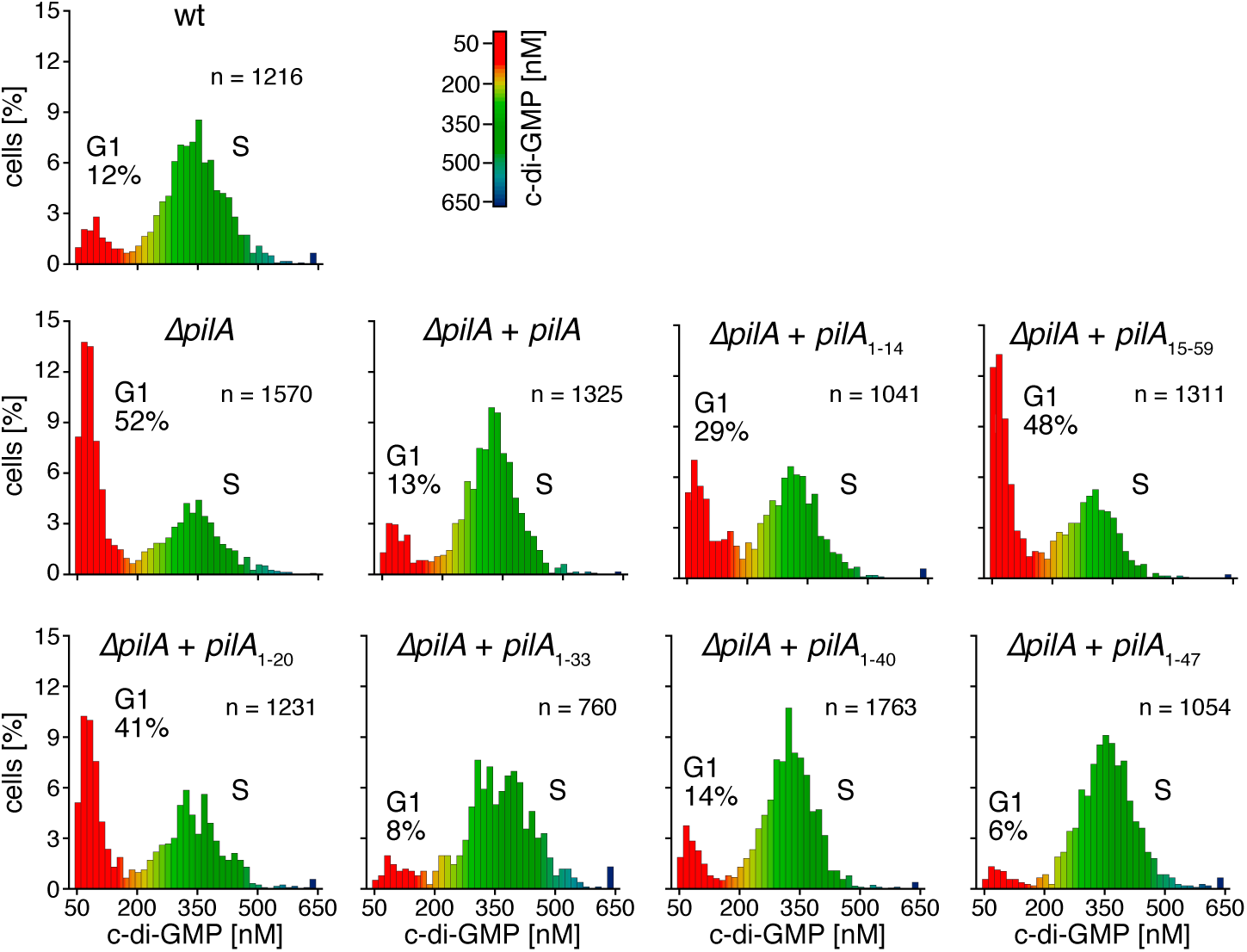
Population distribution of intra-cellular c-di-GMP concentrations in synchronized *Caulobacter* cells with different *pilA* backgrounds assessed by FRET microscopy. Cells with low c-di-GMP levels correspond to swarmer cells (G1) while cells with high c-di-GMP concentrations correspond to replication competent stalked cells (S).

**Fig. S6.**
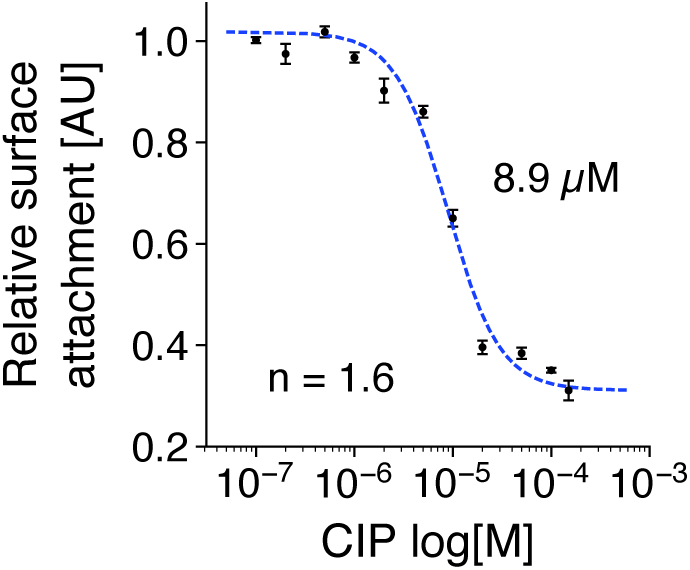
Concentration response curve of the CIP peptide inhibiting cellular attachment with a EC50 of 8.9 and a hill coefficient of 1.6.

**Table S1.** Minimal generation times of *Caulobacter* in PYE medium at selected growth temperatures.

| Growth temperature [°C] | Generation time [h] |
| --- | --- |
| 5 | 52.6<sup>a</sup> |
| 10 | 18.2 ± 0.24 |
| 25 | 1.7 ± 0.04 |
| 30 | 1.4 ± 0.02 |

<sup>a</sup> Calculated generation time according to Ratkowsky form.

**Table S2.** Transposon sequencing statistics.

| **Tn5 mutant library** | **Mapped reads<sup>a</sup>** | **Unique Tn5 insertions<sup>b</sup>** | **Mean gap [bp]<sup>c</sup>** | **Median gap [bp]<sup>c</sup>** |
| --- | --- | --- | --- | --- |
| 5 °C | 7'838'227 | 398'150 | 10 | 5 |
| 10 °C | 9'726'083 | 502'774 | 8 | 4 |
| 25 °C | 9'773'590 | 397'377 | 10 | 5 |
| 30 °C | 10'380'804 | 399'751 | 10 | 5 |

<sup>a</sup> Number of unambiguously mapped paired-end reads.
<sup>b</sup> Number of unique transposon insertions mapped.
<sup>c</sup> Gap between consecutive transposon insertions.

**Table S3.** Complementation analysis of PilA variants and distribution of cell cycle states according to single cell c-di-GMP measurement.

| Genotype | G1 phase<br>[n cells] | S phase<br>[n cells] | Fraction in<br>G1 phase [%] | P-value<sup>a</sup> |
| --- | --- | --- | --- | --- |
| wt | 143 | 1073 | 11.76 | - |
| $\Delta pilA$ | 821 | 749 | 52.29 | $4.96 \cdot 10^{-119}$ |
| $\Delta pilA + pilA$ | 174 | 1151 | 13.13 | $3.07 \cdot 10^{-1}$ |
| $\Delta pilA + pilA_{1-14}$ | 304 | 737 | 29.20 | $3.60 \cdot 10^{-25}$ |
| $\Delta pilA + pilA_{15-59}$ | 623 | 688 | 47.52 | $5.83 \cdot 10^{-90}$ |
| $\Delta pilA + pilA_{1-20}$ | 499 | 732 | 40.54 | $2.71 \cdot 10^{-61}$ |
| $\Delta pilA + pilA_{1-33}$ | 62 | 698 | 8.16 | $1.20 \cdot 10^{-3}$ |
| $\Delta pilA + pilA_{1-40}$ | 241 | 1522 | 13.67 | $1.33 \cdot 10^{-1}$ |
| $\Delta pilA + pilA_{1-47}$ | 59 | 995 | 5.60 | $1.20 \cdot 10^{-7}$ |

<sup>a</sup> Fisher's exact test of observing the measured population distribution under the null hypothesis that cell cycle states are distributed as in the wt.

**Table S4.** Essentiality of the pilus assembly genes within the Tad locus.

| Locus_tag | Gene | Functional annotation | Essentiality |
| --- | --- | --- | --- |
| CCNA_03043 | *pilA* | type IVb pilin protein | essential for slow growth |
| CCNA_03042 | *cpaA* | Prepilin peptidase | non-essential |
| CCNA_03041 | *cpaB* | Periplasmic pilus assembly subunit | non-essential |
| CCNA_03040 | *cpaC* | Outer membrane pilus secretion channel | non-essential |
| CCNA_03039 | *cpaD* | Periplasmic pilus assembly subunit | non-essential |
| CCNA_03038 | *cpaE* | Pilus assembly ATPase | non-essential |
| CCNA_03037 | *cpaF* | Extension and Retraction ATPase | non-essential |
| CCNA_03036 | *cpaG* | TadB-related pilus assembly protein | non-essential |
| CCNA_03035 | *cpaH* | TadC-related pilus assembly protein | non-essential |
| CCNA_03044 | *cpaI* | CpaC-related secretion pathway protein | non-essential |
| CCNA_03045 | *cpaJ* | Pseudopilin | non-essential |
| CCNA_03046 | *cpaK* | Pseudopilin | non-essential |

